## Supplementary material for "Elevated exposure to prenatal thyroid hormones affects embryonic mortality but has no clear effects into adulthood"

### Effects of prenatal THs on early development

##### Duration of embryonic period

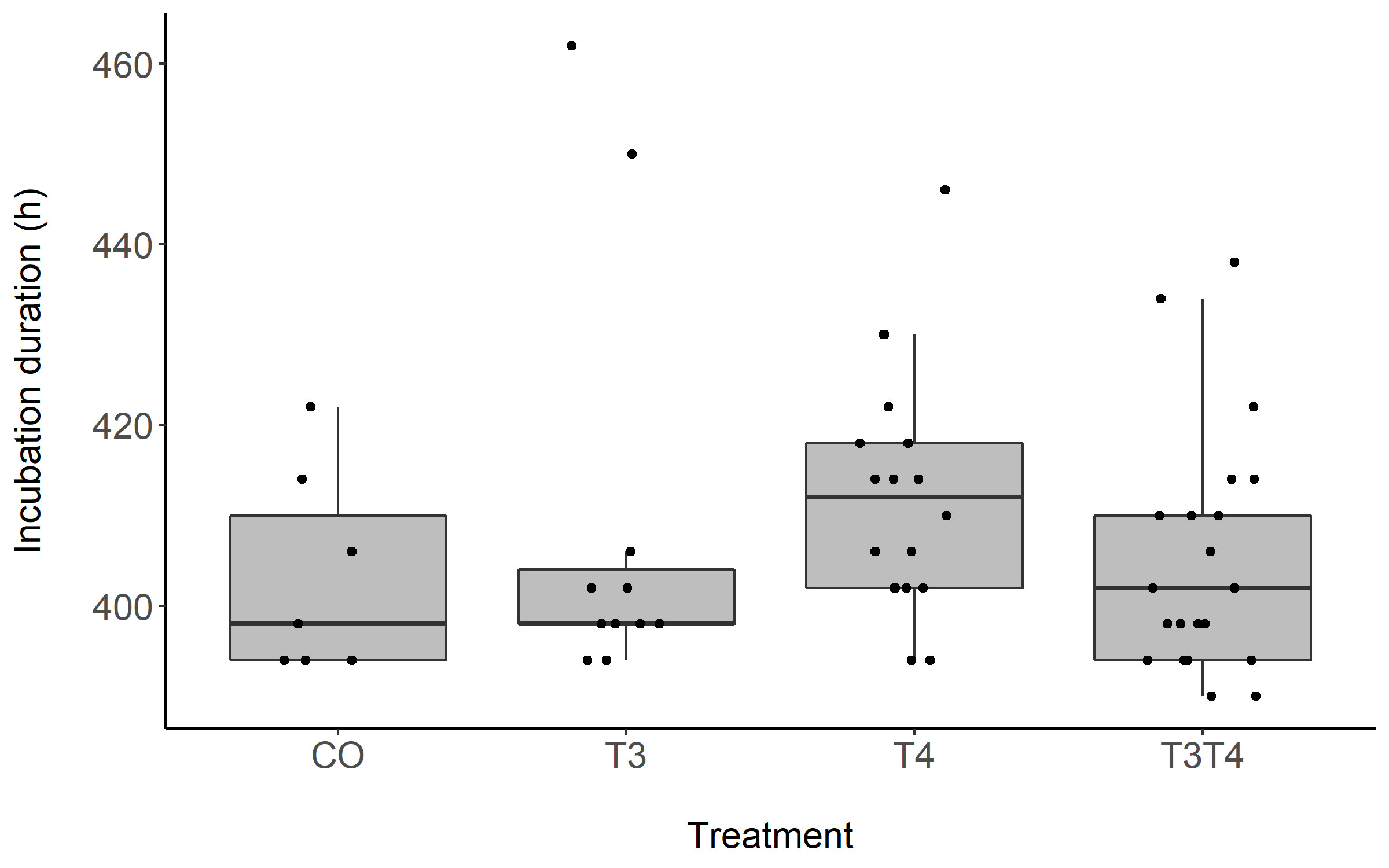

Figure S1: Boxplot of the duration of the embryonic period in Japanese quail eggs according to yolk TH manipulation treatments: CO (N=10), T_3_ (N=15), T_4_ (N=20), T_3_T_4_ (N=21). CO = control, T_4_ (thyroxine) = injection of T_4_, T_3_ (triiodothyronine) = injection of T_3_, T_3_T_4_ = injection of T_3_ and T_4_.

##### Mass at hatching

Figure S2: Boxplot of the mass at hatching in Japanese quail chicks according to yolk TH manipulation treatments: CO (N=10), T_3_ (N=15), T_4_ (N=20), T_3_T_4_ (N=21). See Fig. S1 for a description of the treatments.

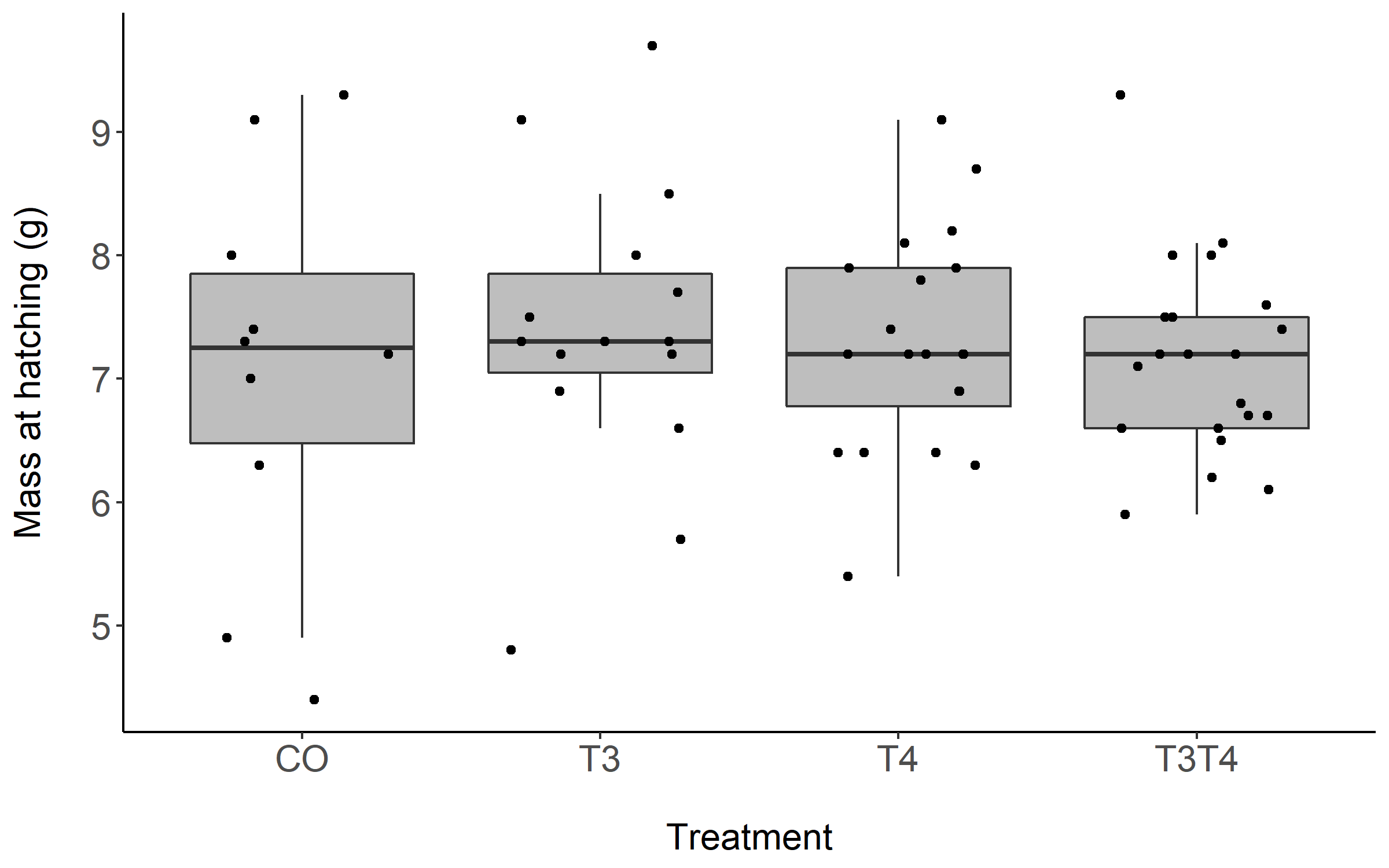

#### Effects of prenatal THs on tarsus and wing growth

Table S1: Results of the Generalised Additive Mixed Models (GAMMs) on wing length, with sex, treatment and their interaction fitted either as intercept, curve shape or both (all combinations tested). A total of 12 GAMMs were fitted and ranked based on their AIC, from the lowest to the highest. Weight: Akaike’s weights. ER: the evidence ratio of the weight of the top-supported model divided by the weight of the null model (model 12).

| Model | Intercept | Curve shape | ΔAIC | df | Weight | ER |
| --- | --- | --- | --- | --- | --- | --- |
| 11 | Sex | - | 0.0 | 9 | 0.60 | 4 |
| 9 | Treatment + Sex | - | 2.2 | 12 | 0.21 | - |
| 12 | - | - | 2.8 | 8 | 0.15 | - |
| 10 | Treatment | - | 5.4 | 11 | 0.04 | - |
| 1 | Sex | Sex | 29.3 | 11 | <0.001 | - |
| 3 | - | Sex | 30.0 | 10 | <0.001 | - |
| 8 | Treatment + Sex | Sex | 31.2 | 14 | <0.001 | - |
| 2 | Treatment | Sex | 32.4 | 13 | <0.001 | - |
| 5 | Sex | Treatment | 102.6 | 15 | <0.001 | - |
| 7 | Treatment + Sex | Treatment | 104.1 | 18 | <0.001 | - |
| 6 | - | Treatment | 105.1 | 14 | <0.001 | - |
| 4 | Treatment | Treatment | 107.4 | 17 | <0.001 | - |

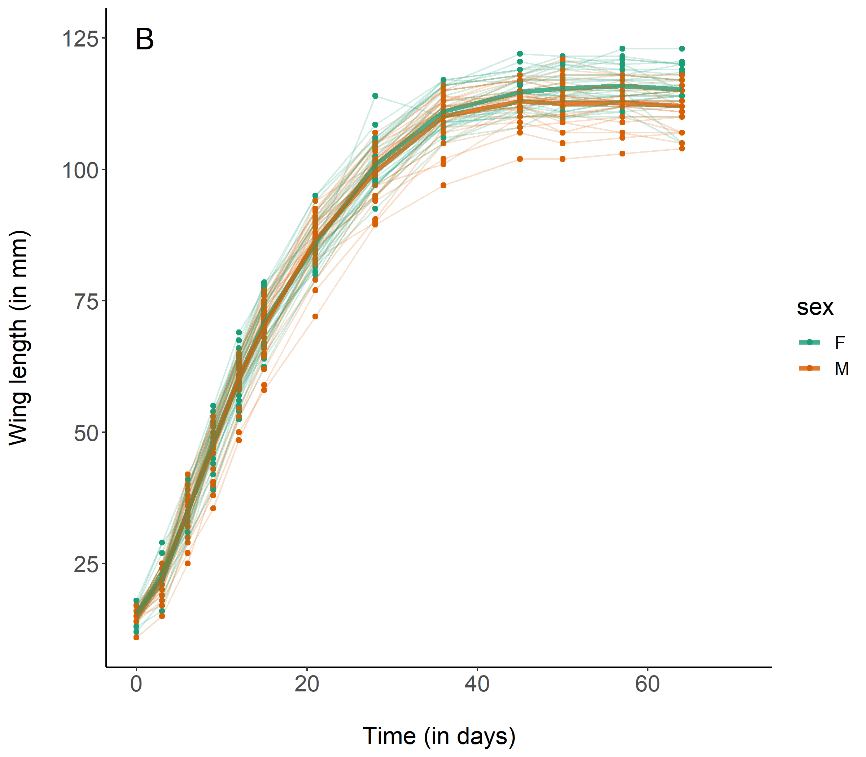

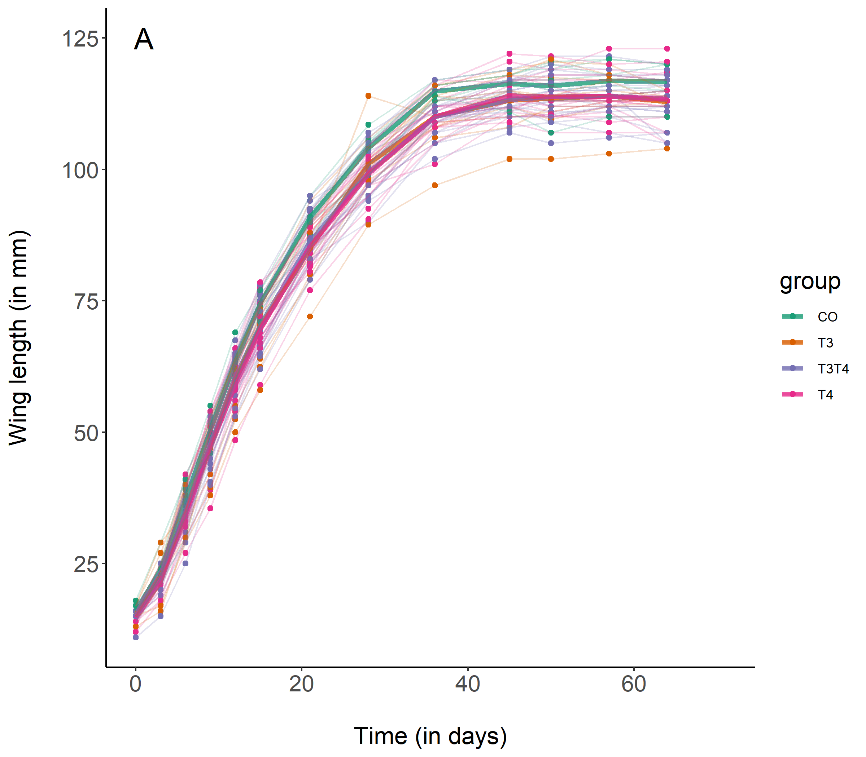

Figure S3: Growth curves in wing length of Japanese quails hatching from eggs treated with either T_3_, T_4_, a combination of both hormones, or a control solution. See Fig. S1 for a description of the treatments. Each line represents an individual bird, while thick coloured lines represent mean values. A: Growth curve according to yolk TH manipulation. N = 7 CO, 11 T_3_, 18 T_4_ and 21 T_3_T_4_. B: Growth curve according to sex. N = 29 females and 28 males.

Table S2: Results of the Generalised Additive Mixed Models (GAMMs) on tarsus length, with sex, treatment and their interaction fitted either as intercept, curve shape or both (all combinations tested). A total of 12 GAMMs were fitted and ranked based on their AIC, from the lowest to the highest. Weight: Akaike’s weights. ER: the evidence ratio of the weight of the models within ΔAIC ≤ 2 divided by the weight of the null model (model 12).

| Model | Intercept | Curve shape | ΔAIC | df | Weight | ER |
| --- | --- | --- | --- | --- | --- | --- |
| 10 | Treatment | - | 0.0 | 11 | 0.46 | 3.5 |
| 9 | Treatment + Sex | - | 1.0 | 12 | 0.28 | 2.2 |
| 12 | - | - | 2.5 | 8 | 0.13 | - |
| 11 | Sex | - | 2.5 | 9 | 0.13 | - |
| 2 | Treatment | Sex | 21.4 | 13 | <0.001 | - |
| 8 | Treatment + Sex | Sex | 22.8 | 14 | <0.001 | - |
| 3 | - | Sex | 23.5 | 10 | <0.001 | - |
| 1 | Sex | Sex | 24.2 | 11 | <0.001 | - |
| 4 | Treatment | Treatment | 91.3 | 17 | <0.001 | - |
| 7 | Treatment + Sex | Treatment | 92.4 | 18 | <0.001 | - |
| 6 | - | Treatment | 943.8 | 14 | <0.001 | - |
| 5 | Sex | Treatment | 94.1 | 15 | <0.001 | - |

Figure S4: Growth curves in tarsus length of Japanese quails hatching from eggs treated with either T_3_, T_4_, a combination of both hormones, or a control solution. See Fig. S1 for a description of the treatments. Each line represents an individual bird, while thick coloured lines represent mean values. A: Growth curve according to yolk TH manipulation. N = 7 CO, 11 T_3_, 18 T_4_ and 21 T_3_T_4_. B: Growth curve according to sex. N = 29 females and 28 males.

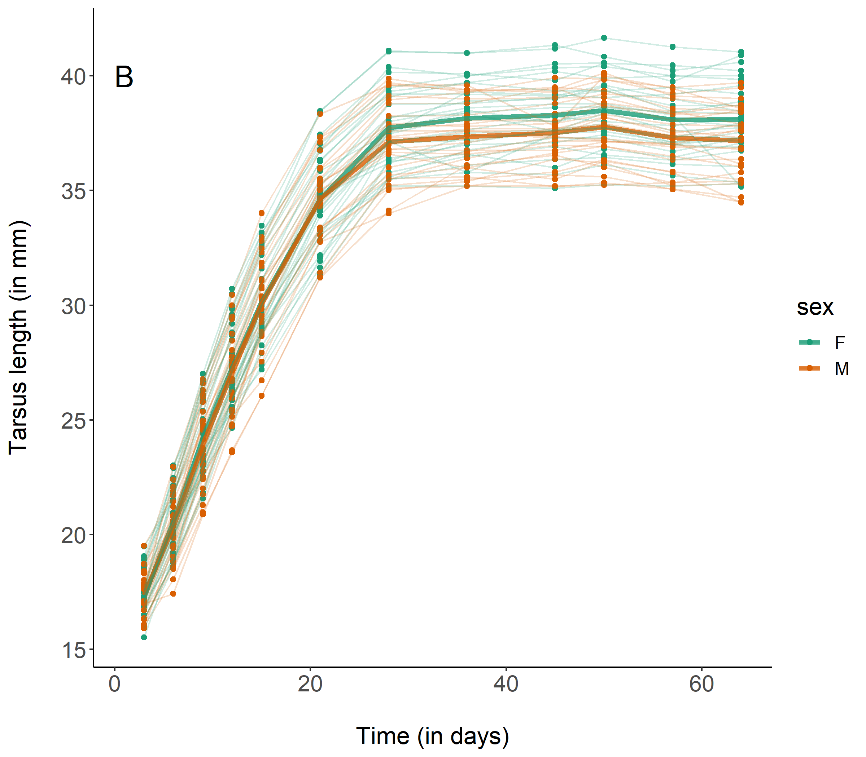

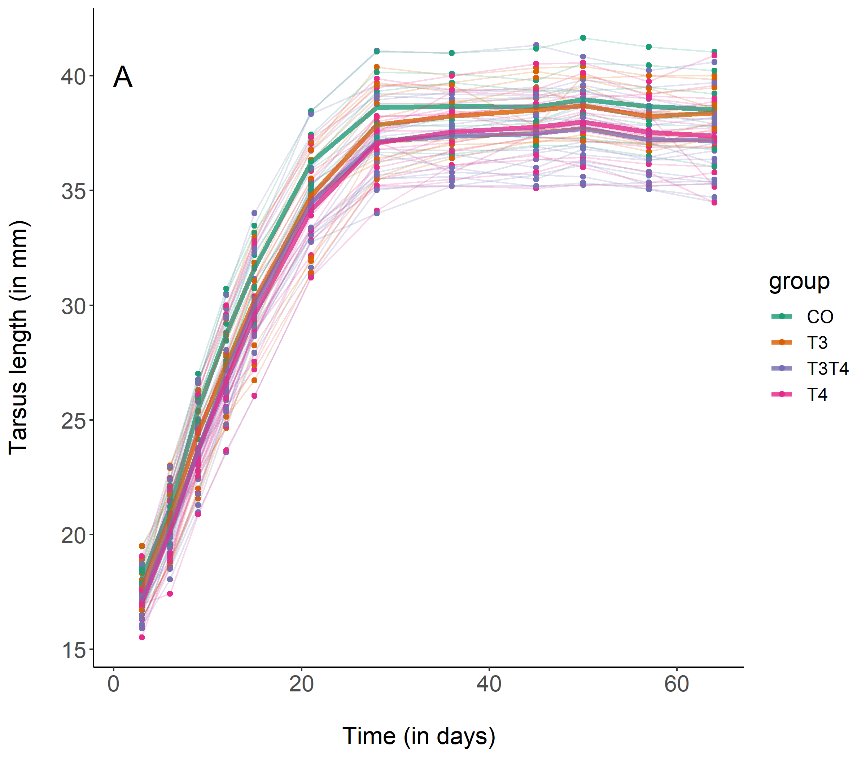

##### **Reproductive maturation and regression, and female reproductive investment**

*Statistical analyses*

Male reproductive maturation was assessed by measuring the growth of the cloacal gland and foam production from week 4 to week 10 after hatching. Cloacal gland growth is non-linear and was then fitted with a GAMM with the intercept and the curve shape varying according to the treatment, similarly as for body mass. The model included a residual autocorrelation structure AR-1 to account for repeated measurements (Zuur, 2009).

Foam production was fitted with an LMMs with treatment as a predictor and body mass as a covariate. Two- and three-way interactions between these three parameters were tested, but not presented as statistically non-significant. Mother identity was added as a random intercept, and individuals were nested within mothers and allowed to vary both in the intercept and in the slope (i.e., random slope).

Female reproductive investment in eggs was assessed by multiplying the mean egg mass by the number of eggs laid by 2-month-old females over 6 days (N = 27). Data were analysed by separate LMMs with treatment as the fixed factor and mother identity of the parental generation as a random intercept. Female body mass two days before egg collection was added as a covariate.

Cloacal gland regression in males was analysed with an identical LMM as for foam production.

#### Effects of prenatal THs on prenuptial life stage transition and reproductive investment

Cloacal gland growth did not differ according to the treatment, as the null model was the only one within ΔAIC ≤ 2 (Table S3, Fig. S5). Likewise, the elevation of yolk THs did not affect foam production (LMM, *F*_3,17.2_ = 0.82, p = 0.50, Fig. S6), while body mass had a positive association with foam production (LMM, Estimate *=* 0.01±0.003, *F*_1,28.6_ = 10.6, p = 0.003). Manipulation of prenatal THs had no effect on female reproductive investment in eggs (LMM, *F*_3,14.8_ = 0.30, p = 0.83, Fig. S7). Taken together, male gonadal development and female reproductive investment were not affected by elevated yolk THs. However, the low sample sizes in the control groups do not allow us to make robust comparisons.

Table S3: Results of the Generalised Additive Mixed Models (GAMMs) on cloacal gland growth, with treatment fitted either as intercept, curve shape or both. A total of 4 GAMMs were fitted and ranked based on their AIC, from the lowest to the highest. Weight: Akaike’s weights.

| Model | Intercept | Curve shape | ΔAIC | df | Weight |
| --- | --- | --- | --- | --- | --- |
| 4 | - | - | 0.0 | 7 | 1 |
| 3 | - | Treatment | 31.2 | 13 | <0.001 |
| 1 | Treatment | Treatment | 35.1 | 16 | <0.001 |
| 2 | Treatment | - | 71.0 | 10 | <0.001 |

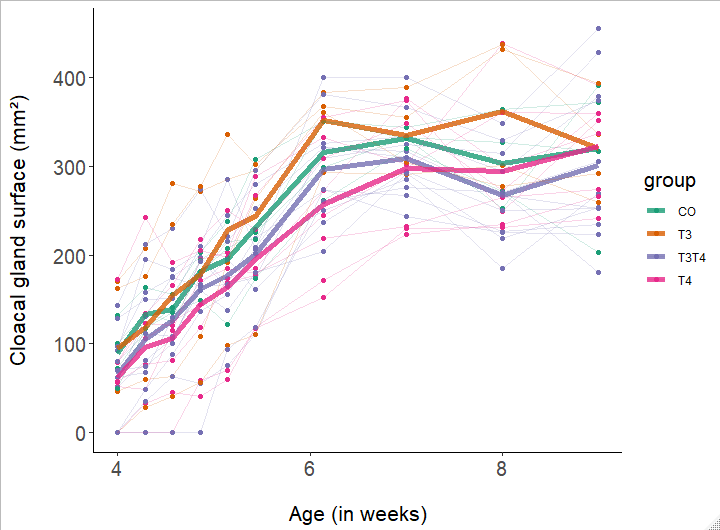

Figure S5: Cloacal gland growth in male Japanese quails according to yolk TH manipulation treatments: CO (N = 4), T_3_ (N = 4), T_4_ (N = 8), T_3_T_4_ (N = 12). See Fig. S1 for a description of the treatments. Measures were taken every second day from week 4 to week 6 after hatching, and then once a week until week 10. Each line represents an individual bird, while thick coloured lines represent mean values.

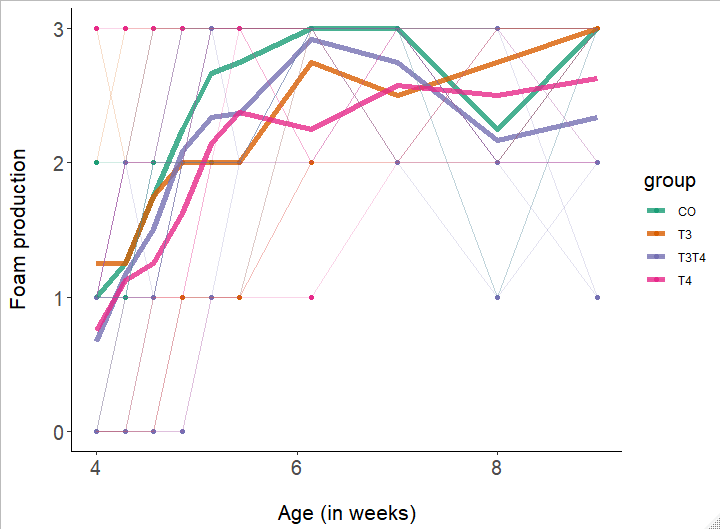

Figure S6: Foam production in male Japanese quails according to yolk TH manipulation treatments: CO (N=4), T_3_ (N=4), T_4_ (N=8), T_3_T_4_ (N=12). See Fig. S1 for a description of the treatments. Measures were taken every second day from week 4 to week 6 after hatching, and then once a week until week 10. Each line represents an individual bird, while thick coloured lines represent mean values.

Figure S7: Boxplot of the reproductive investment by 2-month old female Japanese quails according to yolk TH manipulation treatments: CO (N=3), T_3_ (N=7), T_4_ (N=8), T_3_T_4_ (N=8). Egg investment is calculated as the number of eggs produced during 6 days multiplied by the average egg mass. See Fig. S1 for a description of the treatments.

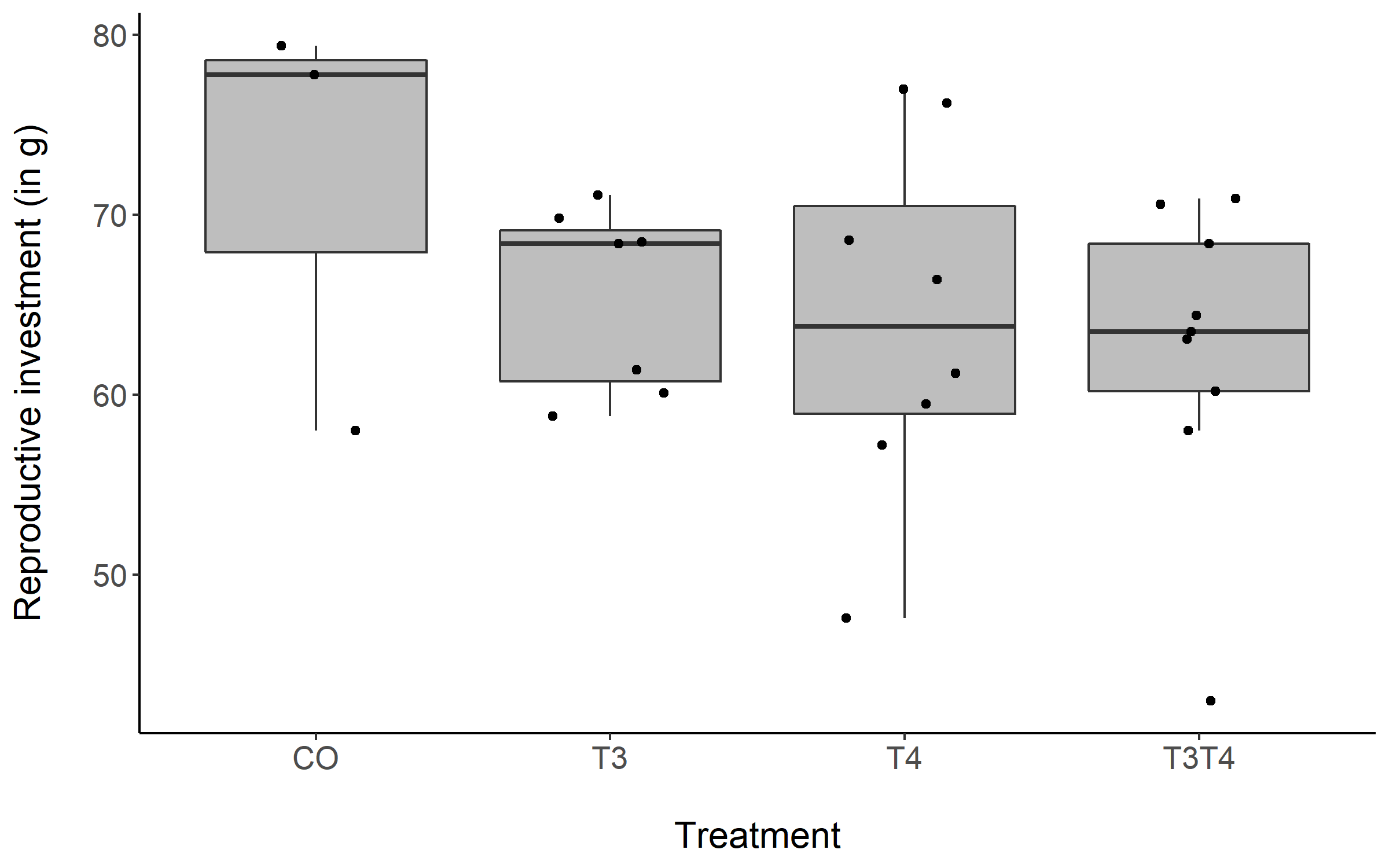

#### Effects of prenatal THs on gonadal regression

Figure S8: Cloacal gland regression of 7-month-old male Japanese quails according to yolk TH manipulation treatments: CO (N = 4), T_3_ (N = 3), T_4_ (N = 7), T_3_T_4_ (N = 12). See Fig. S1 for a description of the treatments. Measures were taken every second day after switching from long photoperiod (16L:8D) to short photoperiod (8L:16D, switch = time point 0 on x-axis). Each line represents an individual bird, while thick coloured lines represent group mean values.

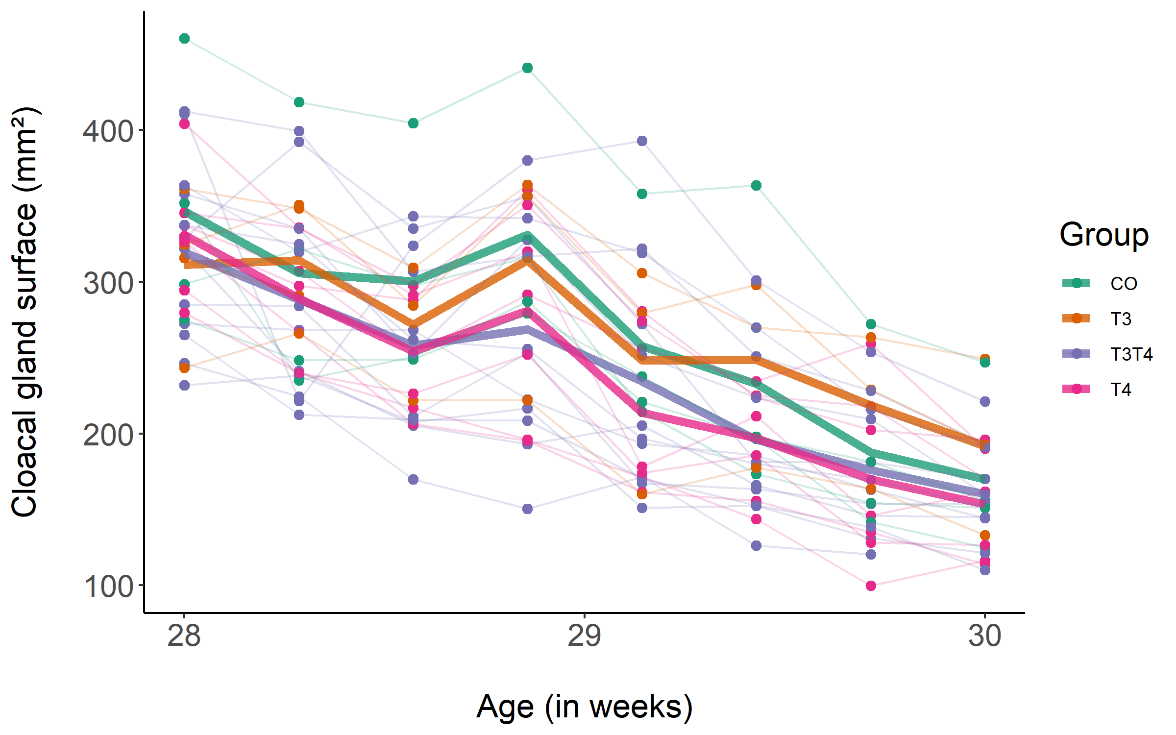

As expected, cloacal gland regressed rapidly after switching to short photoperiod (LMMs, Estimate±SE = -11.61±0.47 mm², *F*_1,183.5_ = 637.58, p < 0.0001, Fig. S8). We detected no effect of yolk thyroid hormones on cloacal gland regression (LMM interaction treatment × time, *F*_3,183.3_ = 1.76, p = 0.16; treatment *F*_3,15.9_ = 0.58, p = 0.64, Fig. S8), while we found a significant positive correlation with body mass (LMM, Estimate±SE = 1.32±0.26 mm², *F*_1,53.2_ = 18.20, p < 0.0001). To sum up, male gonadal regression was not affected by elevated yolk THs. However, the low sample size in the control group does not allow us to make robust comparisons.
